# Early embryogenesis and organogenesis in the annelid *Owenia fusiformis*

**DOI:** 10.1101/2021.02.20.431505

**Authors:** Allan Martín Carrillo-Baltodano, Océane Seudre, Kero Guynes, José María Martín-Durán

## Abstract

**Background:** Annelids are a diverse group of segmented worms within Spiralia, whose embryos exhibit spiral cleavage and a variety of larval forms. While most modern embryological studies focus on species with unequal spiral cleavage nested in Pleistoannelida (Sedentaria + Errantia), a few recent studies looked into *Owenia fusiformis*, a member of the sister group to all remaining annelids and thus a key lineage to understand annelid and spiralian evolution and development. However, the timing of early cleavage and detailed morphogenetic events leading to the formation of the idiosyncratic mitraria larva of *O. fusiformis* remain largely unexplored.

**Results:** *O. fusiformis* undergoes equal spiral cleavage where the first quartet of animal micromeres are slightly larger than the vegetal macromeres. Cleavage results in a coeloblastula approximately five hours post fertilization (hpf) at 19 °C. Gastrulation occurs via invagination and completes four hours later, with putative mesodermal precursors and the chaetoblasts appearing 10 hpf at the dorsoposterior side. Soon after, at 11 hpf, the apical tuft emerges, followed by the first neurons (as revealed by the expression of *elav1* and *synaptotagmin1*) in the apical organ and the prototroch by 13 hpf. Muscles connecting the chaetal sac to various larval tissues develop around 18 hpf and by the time the mitraria is fully formed at 22 hpf, there are FMRFamide^+^ neurons in the apical organ and prototroch, the latter forming a prototrochal ring. As the mitraria feeds, it grows in size and the prototroch expands through active proliferation. The larva becomes competent after ∼3 weeks post fertilization at 15 °C, when a conspicuous juvenile rudiment has formed ventrally.

**Conclusions:** *O. fusiformis* embryogenesis is similar to that of other equal spiral cleaving annelids, supporting that equal cleavage is associated with the formation of a coeloblastula, gastrulation via invagination, and a feeding trochophore-like larva in Annelida. The nervous system of the mitraria larva forms earlier and is more complex than previously recognised and develops from anterior to posterior, which is likely an ancestral condition to Annelida. Altogether, our study identifies the major developmental events during *O. fusiformis* ontogeny, defining a conceptual framework for future investigations.

## Background

Annelids are a diverse and abundant group of segmented worms, part of the larger clade of bilaterian animals called Spiralia (Laumer et al., 2015; Marlétaz et al., 2019). Ancestral to annelids is the presence of the quartet spiral cleavage program during early embryonic development, in which blastomeres divide obliquely and perpendicular to the animal-vegetal axis from the 4-cell stage onwards, alternating directions and giving as a result the stereotypical spiral-like arrangement of the embryonic cells when looked from above (Brun-Usan et al., 2017; Hejnol, 2010; Henry, 2014; Martin-Duran and Marletaz, 2020; Seaver, 2014). Most modern molecular studies of annelid embryogenesis have focused on species belonging to the two most diverse groups, namely Sedentaria (e.g. the capitellid *Capitella teleta*, the leech *Helobdella robusta* and the serpulids *Hydroides elegans* and *Spirobranchus lamarckii*) and Errantia (e.g. the nereidid *Platynereis dumerilii*), which together form Pleistoannelida (Bleidorn et al., 2015; Seaver, 2014; Shankland and Seaver, 2000; Williams and Jékely, 2016) (Fig. 1a). However, the current annelid phylogeny shows Pleistoannelida as a deeply nested clade, with up to five intermediate lineages between this group and the last common annelid ancestor (Helm et al., 2018; Martín-Durán et al., 2020; Weigert et al., 2014) (Fig. 1a). Therefore, investigating these other early branching lineages, and in particular Palaeoannelida (Oweniidae + Magelonidae) as the sister taxon to all other annelids (Fig. 1a), is fundamental to uncover the origins and early diversification of Annelida (Helm et al., 2016; Helm et al., 2018; Martín-Durán et al., 2018).

**Figure 1.**
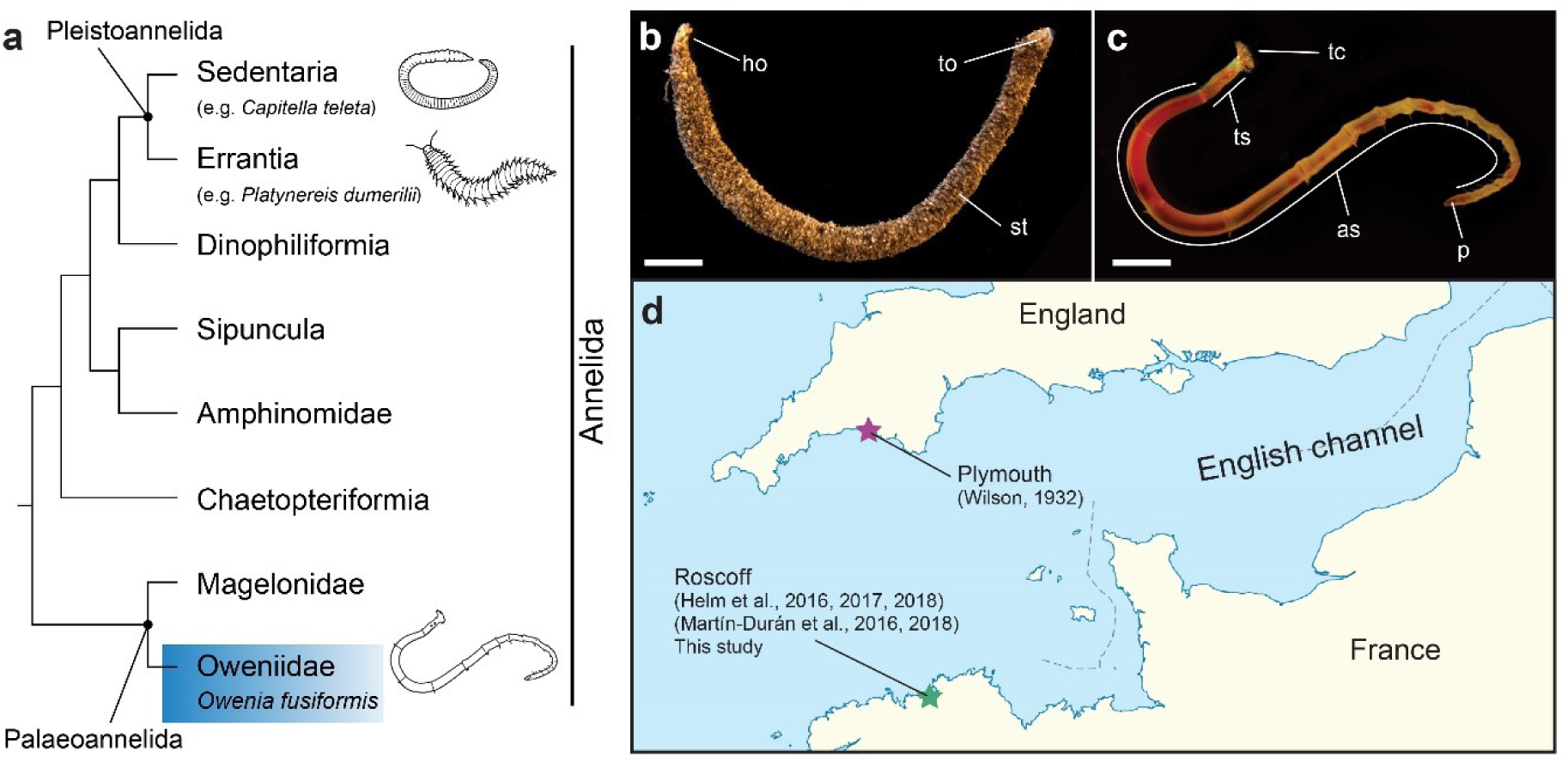
*Owenia fusiformis*, phylogenetic position and sampling location. (**a**) *O. fusiformis* (blue box) is a member of Palaeoannelida, the sister clade to all remaining annelids. Annelid phylogeny according to (Martín-Durán et al., 2020); (**b, c**) As adult, *O. fusiformis* dwells inside a self-built sand tube. The adult body is divided into a head with tentacles, three thoracic segments, abdominal segments and a pygidium (**c**); (**d**) Embryological studies on *O. fusiformis* have focused on specimens collected from the English channel, with the most recent ones studying a population near the Marine Biological Station of Roscoff, France. as: abdominal segments; ho: head opening; p: pygidium; st: sand tube; tc: tentacles; to: tail opening; ts: thoracic segments. Scale bar is 2 cm.

Embryological studies in Palaeoannelida are however rare and have mostly centered on the genus *Owenia*. Wilson (Wilson, 1932) focused on morphological descriptions of the idiosyncratic mitraria larva up to metamorphosis in *Owenia fusiformis* Delle Chiaje, 1844, without providing much information about embryogenesis. Early development has only been described for *Owenia collaris* Hartman, 1955 from the North West Pacific (Smart and Von Dassow, 2009), which exhibits equal spiral cleavage and gastrulation via invagination. Later studies on the serotonergic and FMRFamidonergic nervous system were carried out in *O. fusiformis* (Helm et al., 2016), preliminarily describing the larval nervous system as a complex of a few neurons and axons that develop relative late in the mitraria (∼48 hours post fertilization). Recently, a set of studies analyzing the expression of the genetic toolkit responsible of axial identity and germ layer specification established *O. fusiformis* as the reference and more tractable research species for molecular investigations within Palaeoannelida (Martín-Durán et al., 2016a; Martín-Durán et al., 2016b; Martín-Durán et al., 2018) (Fig. 1b–d). However, the early development of *O. fusiformis* has not been described yet, and a systematic and detailed description of the major morphogenetic events after gastrulation and during larval development is lacking.

Here, we combine confocal laser-scanning microscopy and immunostainings, with cell proliferation assays and gene expression analyses to provide a high-resolution characterization of the embryonic development and larval growth of *O. fusiformis* (Fig. 1b–d). We identify the onset of major developmental events in *O. fusiformis*, describing cleavage, gastrulation and organogenesis (myogenesis, neurogenesis, chaetogenesis and ciliogenesis) and establishing morphological and molecular landmarks that define a consistent staging system for this annelid species. Altogether, our study contributes to a more systematic and comprehensive understanding of the embryonic development of *O. fusiformis* and Palaeoannelida generally, which proves essential to better reconstruct ancestral developmental traits to Annelida and Spiralia.

## Results

### Spiral cleavage

Females of *O. fusiformis* spawn small oocytes (∼100µm) that are flat and exhibit a conspicuous germinal vesicle (Fig. 2a). Oocyte activation occurs naturally in sea water, resulting in the germinal vesicle breaking down and the oocytes becoming more spherical and receptive to sperm. After fertilization, the zygote (Fig. 2b) takes about 30–60 min at 19 °C to extrude the polar bodies and undergo the first holoblastic cleavage, which produces two blastomeres of equal size (Fig. 2c). Within 30 min, the 2-cell embryo divides again symmetrically and dextrally, and by 1.5 hours post fertilization (hpf) the second zygotic division generates four equal blastomeres, two of which share a vegetal cross furrow (Fig. 2d–f). Half an hour later (∼2 hpf), the 4-cell stage embryo divides dextrally and perpendicular to the animal-vegetal axis to form the first quartet of animal micromeres (1q) and four vegetal macromeres (1Q) (Fig. 2g–i). Different to most other annelids, but similar to *O. collaris* (Smart and Von Dassow, 2009), sipunculans (Boyle and Rice, 2014) and nemerteans (Henry and Martindale, 1998; Martindale and Henry, 1995), 1q is slightly larger than the vegetal macromeres (Fig. 2i). Rapidly, the 8-cells stage embryo cleaves sinistrally, giving rise to the second micromere quartet (2q), and the 1q^1^ and 1q^2^ tiers of micromeres (1q^1^ being bigger than 1q^2^), which end up positioned in a similar horizontal plane relative to each other (Fig. 2j–l). This cleavage pattern of the 1q micromeres generates an incipient blastocoel (Fig. 2l). Between 3 and 4 hpf, the fifth, dextral cleavage forms the third quartet of micromeres (3q) and a 32-cell stage embryo (Fig. 2m–n). Five hours after fertilization, the last, sinistral round of cell divisions generates the fourth quartet of micromeres (4q) and the 4Q macromeres (Fig. 2o–p). This 64-cell stage embryo, or mature coeloblastula, has a prominent central blastocoel (Fig. 2p), with the larger vegetal blastomeres forming the gastral plate that will invaginate during gastrulation. *Owenia fusiformis* embryos thus undergo typical equal spiral cleavage, similar to *O. collaris* (Smart and Von Dassow, 2009), and apparently without signs of asymmetry between equivalent blastomeres of each embryonic quadrant, until the transition between the 32-cell to the 64-cell stage, when one of the 3q micromeres starts dividing first (Fig. 2o).

**Figure 2.**
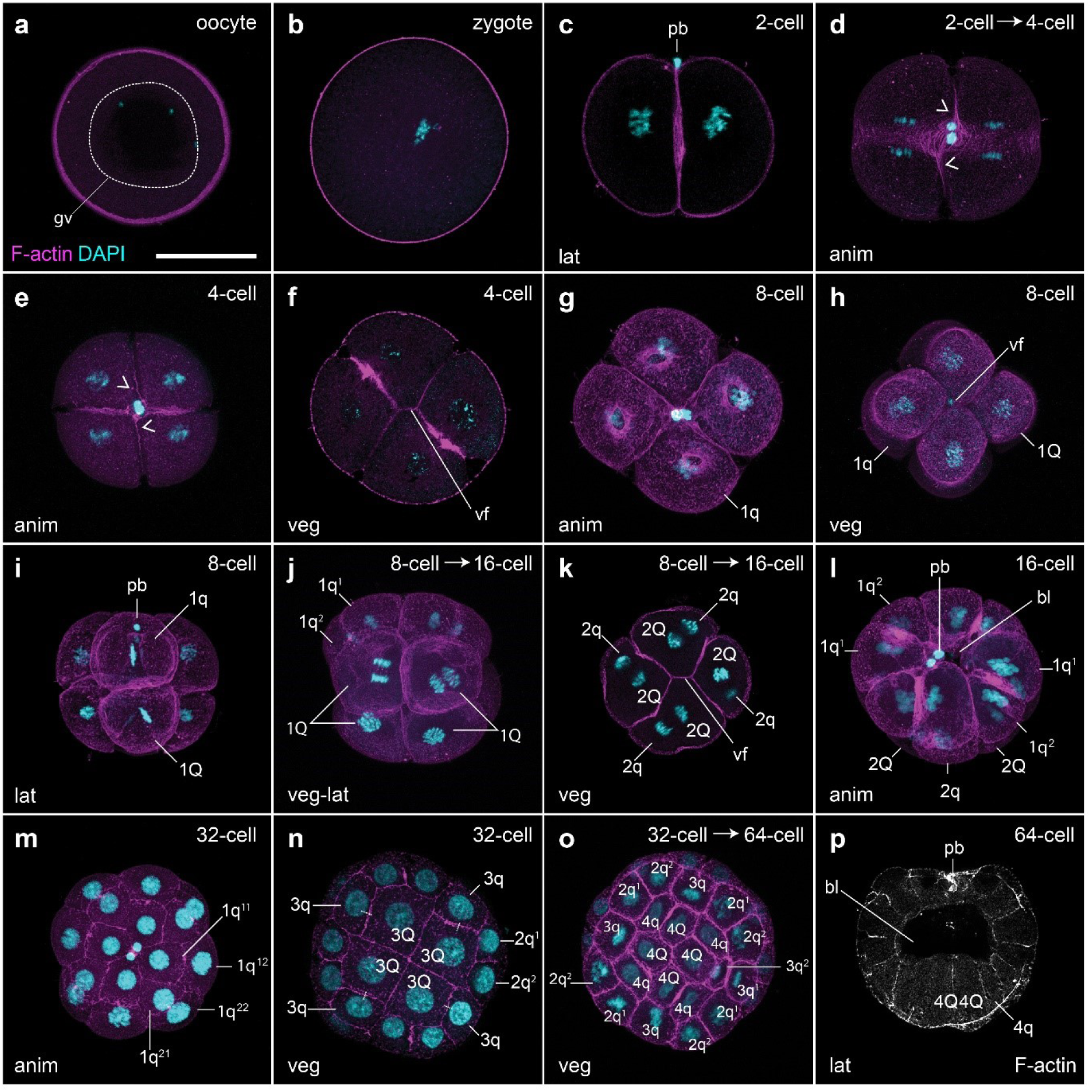
*Owenia fusiformis* undergoes equal spiral cleavage. Confocal laser scanning microscopy (CLSM) images of early embryonic stages from (**a**) oocyte to (**p**) blastula stage (approx. 64-cell). (**a**) The oocyte activates naturally in sea water and after the germinal vesicle breaks down. (**b**) After fertilization, the embryo undergoes equal cleavage resulting in two and later four equal blastomeres (**c–f**). Open arrowheads in (**d**) and (**e**) points to the spiral deformation of the actin cytoskeleton in preparation for the dextral cleavage. (**f–k**) The spiral cleavage initiates with the formation of four oblique micromeres (1q) that are larger than the macromeres (1Q). (**l–o**) Cleavage continues with the stereotypical alternation of the spindles back and forward from counterclockwise to clockwise to give rise to the second (2q) (**l**), third (3q) (**m–n**) and fourth (4q) quartet of micromeres (**o–p**), until the formatin of the coeloblastula (**o**). bl: blastocoel; gv: germinal vesicle; pb: polar bodies; vf: vegetal cross furrow. Scale bar is 50 µm.

### Gastrulation

Gastrulation begins around 5.5–6 hpf via the invagination of the vegetal gastral plate (Fig. 3a), with the 5Q macromeres becoming apically constricted at the vegetal pole and leading the invaginating front, which eventually opens a blastopore by 6 hpf (Fig. 3b). Seven hours after fertilization (7 hpf), the archenteron roof reaches the basal side of the animal ectoderm, making contact with at least one animal cell (Fig. 3c, inset). At this stage, the polar bodies appear internalized, between the archenteron roof and the animal ectoderm (Fig. 3d). Gastrulation completes by 9 hpf (Fig. 3e–f). At this stage, which we refer to as the gastrula stage, the blastocoel is nearly completely obliterated, the archenteron cavity is fully formed, and the blastoporal opening and its rim occupy the entire vegetal pole. At 10 hpf, two mesodermal precursor cells are visible in the blastocoel (Fig. 3g) and there is a band of dividing ectodermal cells located slightly vegetally to the equator (Fig. 3h). We interpret this row of cells as the presumptive prototroch, identified as the 1q^2222^ lineage in *O. collaris* (Smart and Von Dassow, 2009). In addition, at least two chaetoblasts appear on the same side of the vegetal pole as the mesodermal precursors (Fig. 3h). Because the chaetal sac will form on the dorsoposterior part of the embryo (Helm et al., 2016; Martín-Durán et al., 2016a; Smart and Von Dassow, 2009; Wilson, 1932), we deem the region of the gastrula where these chaetoblasts appear as the posterior end. Therefore, its appearance is the first evident morphological landmark revealing the anterior-posterior and dorsal-ventral axes of the embryo and marks the beginning of the axial elongation and organogenesis in the embryos of *O. fusiformis*.

**Figure 3.**
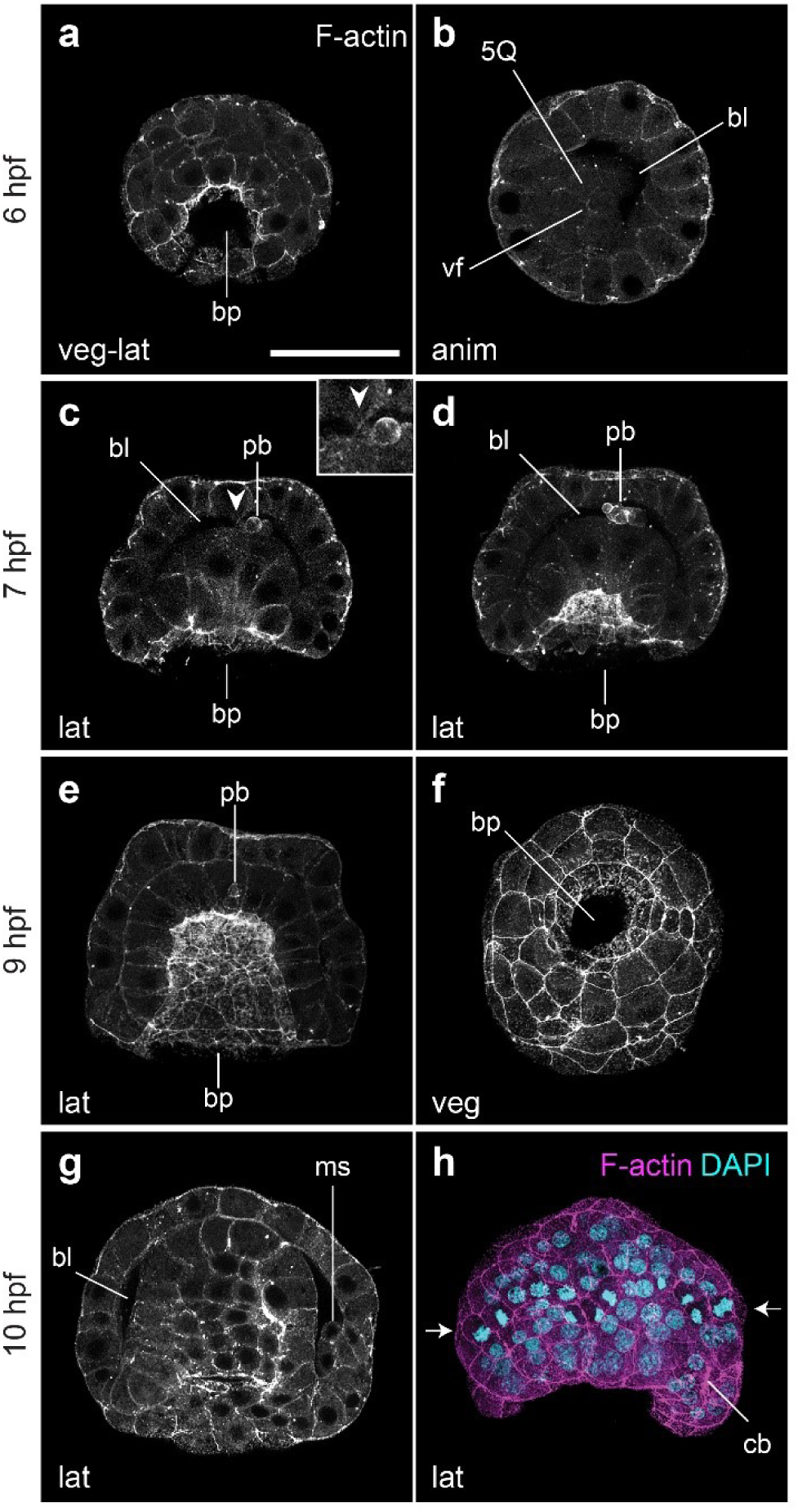
Gastrulation via invagination in *O. fusiformis*. CLSM images of early embryonic stages from (**a**) 6 hpf (beginning of gastrulation) to (**h**) 10 hpf (post gastrulation). (**a**) At 6 hpf the gastral plate starts invaginating with the 5Q macromeres at the front of the archenteron roof (**b**). (**c**) By 7 hpf the archenteron roof connects with the animal ectoderm (closed arrowhead) (inset), and the polar bodies have internalized into the blastocoel (**c–d**). (**e**) Gastrulation ends by 9 hpf, in which the two germinal layers: endoderm and ectoderm are formed; while the mesoderm precursors and chaetoblasts form at the dorso-posterior end of the embryo (**g–h**). Arrows in (**h**) show the row of presumptive prototrochal cells dividing. bl: blastocoel; bp: blastopore; gv: germinal vesicle; ms: mesodermal precursor cells; pb: polar bodies; vf: vegetal cross furrow. Scale bar is 50 µm.

### Organogenesis and development of the early mitraria larva

The first external signs of organogenesis become evident at 11 hpf, when the prototroch, the largest ciliary band and main locomotive structure of the mitraria larva, becomes apparent at a comparable subequatorial ectodermal region with actively dividing cells at 10 hpf (Fig. 3h). Although the prototrochal cells are not yet fully ciliated at 11 hpf, they define two large embryonic regions, namely the episphere on the former animal region and the hyposphere on the original vegetal pole. At this stage in the episphere, the first cilia differentiate at the apical tuft (Fig. 4a–c), as well as seven large cells of unknown function, named refringent globules by Wilson (Wilson, 1932). In the hyposphere, the dorsoposterior chaetal sac forms, bearing four chaetoblasts, and the blastoporal opening narrows and elongates along the anterior-posterior axis (Fig. 4a). At 13 hpf, cells in the mid-posterior and lateral sides of the blastoporal rim start constricting, closing the blastoporal opening (Fig. 4d–e). Four long ciliated cells are now present on each side of the blastopore, slightly behind the closing posterior edge of the blastoporal rim. Due to their bilateral symmetry and their position parallel to the anterior-posterior axis, we deem these cilia as a short neurotroch (Fig. 4d–e). At this early stage, there is proliferation in most regions of the larval body, as we observe DNA replication (EdU signal) and mitotic activity (phospho-Histone H3 signal) especially in the hyposphere and the developing gut, but also in the apical organ and the ciliary bands (Fig. 5a–b).

**Figure 4.**
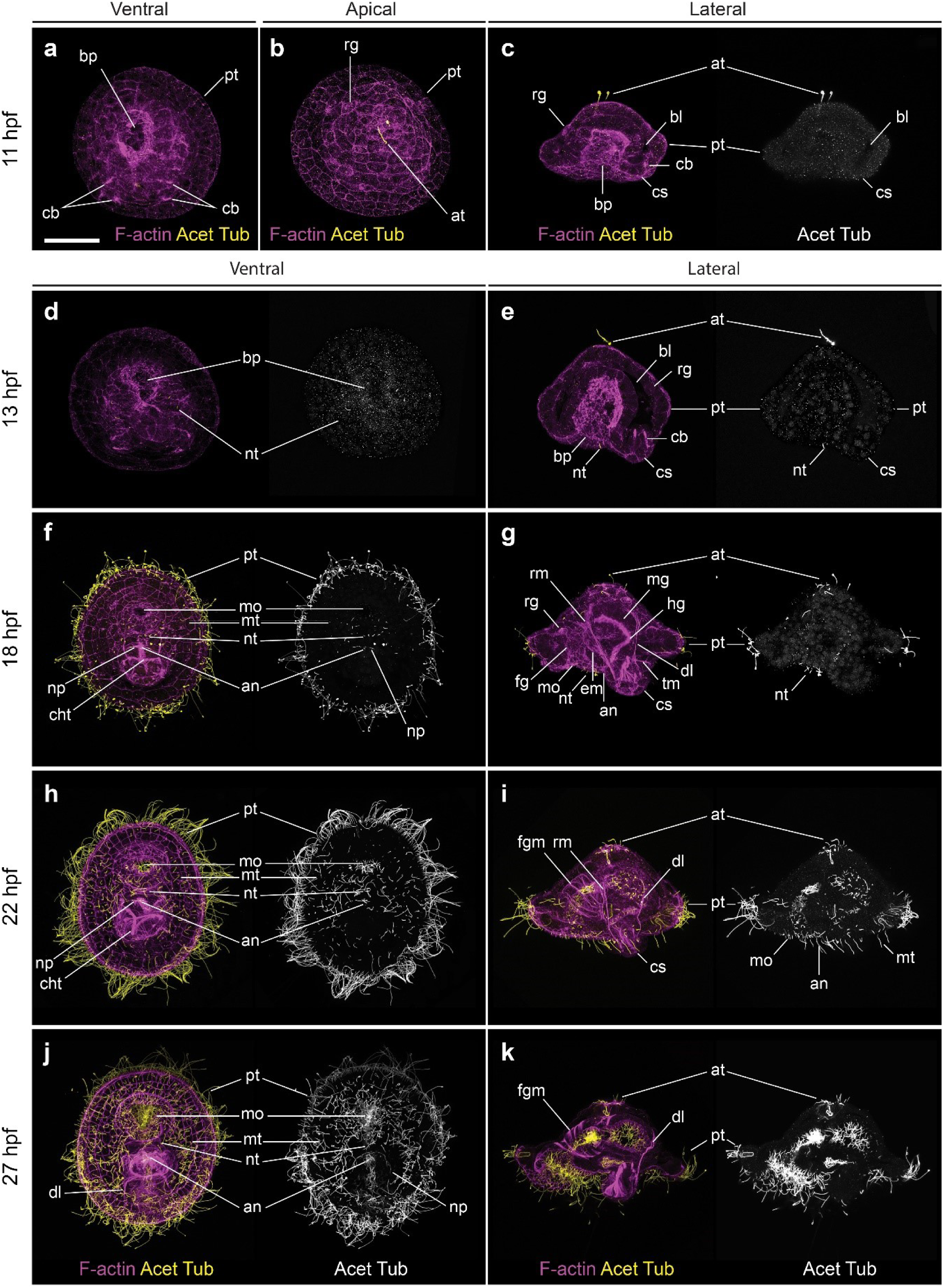
Early mitraria larvae and the beginning of organogenesis. CLSM images of early larval stages from (**a**) 11 hpf to (**k**) 27 hpf. Images on the left column are ventral views with anterior facing up, while those on the right are lateral views with anterior facing left, except for (**b**) which is an apical view. (**a–c**) Ciliogenesis starts at 11 hpf with the formation of the apical tuft and the prototroch dividing the embryo into an apical episphere and a vegetal hyposphere. (**d–e**) A short neurotroch then forms on the posterior side of the blastopore by 13 hpf. By this stage the blastopore elongates to form the presumptive mouth, but still remains open. (**f–g**) By 18 hpf, the early mitraria now has a secondary ciliary band, the metatroch, in addition to a complete gut, larval muscles and chaetae. (**h–k**) Soon after, the larva grows and expands the metatroch throughout the hyposphere, while more muscle develops, including circular muscles around the foregut. an: anus; at: apical tuft; bl: blastocoel; bp: blastopore; cb: chaetoblast; cht: chaetae; cs: chaetal sac; dl: dorsal levator; em: esophagoeal muscle; fg: foregut; fgm: foregut circular muscle; hg: hindgut; mg: midgut; mo: mouth; mt: metatroch; np: nephridia; nt: neurotroch; pt: prototroch; rg: refringent globules; rm: retractor muscle; tm: membrane between chaetal sac and blastocoel. Scale bar is 50 µm.

**Figure 5.**
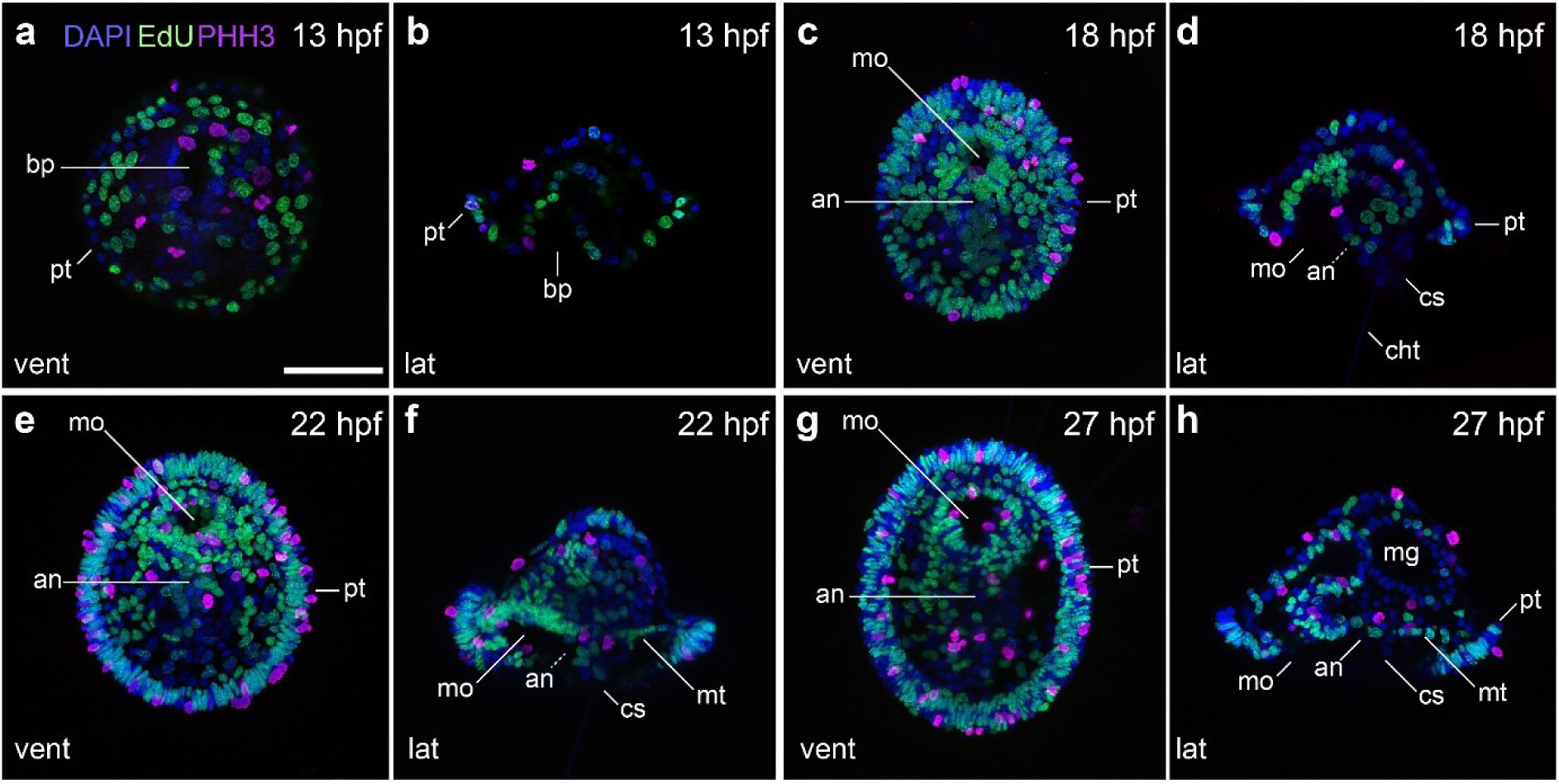
Cell proliferation during early mitraria. CLSM images of EdU and phosphoHistone 3 (PHH3) labeling on early mitraria larvae from (**a**) 13 hpf to (**h**) 27 hpf. (**a**), (**c**), (**e**), (**g**) Are ventral views with anterior facing up, while (**b**), (**d**), (**f**), (**h**) are lateral views with anterior facing left. Cell proliferation is widespread throughout the body, overlapping with cells that are mitotically active in the ciliary bands and the developing gut. an: anus; bp: blastopore; cht: chaetae; cs: chaetal sac; mg: midgut; mo: mouth; mt: metatroch; pt: prototroch. Scale bar is 50 µm.

By 18 hpf, the anterior blastoporal opening has formed the mouth (Martín-Durán et al., 2016a), which is clearly separated from the anus by two or three rows of cells (Fig. 4f) (see also Figure 2 in Wilson (Wilson, 1932)), defining a short ventral side in the embryo. At this stage, the digestive track is divided into foregut, midgut and hindgut, with the neurotroch cells positioned between the mouth and the anus. Importantly, it is at this stage that a new group of scattered ciliated cells starts to emerge in the hyposphere, where the four chaetae now protrude from the chaetal sac (Fig. 5,6). Associated with the enlargement of the prototroch and the appearance of more ciliated cells in the hyposphere, we observe increased proliferation in these areas (Fig. 5). In other trochophore larvae, designated cells called the trochoblasts become arrested and enlarged to form the multiciliated cells of the prototroch (Damen and Dictus, 1994a; Damen and Dictus, 1994b; von Dassow and Maslakova, 2017). However, the mitraria larva has monociliated cells that are able to keep dividing and replacing loss cells within the ciliary bands (Bird, 2012; Smart and Von Dassow, 2009) (Fig. 5). This 18 hpf stage also reveals the first signs of myogenesis, with the appearance of the retractor muscles plus the dorsal levator muscles and esophageal muscles that connect the chaetal sacs to the apical side of the animal and to the esophagus, respectively (Fig. 4g). In addition, two tubulin^+^ nephridia are flanking the anus (Fig. 4e). The increased ciliation at 18 hpf stimulates the swimming behavior of the larva, which starts spinning at the bottom of the culture bowls.

**Figure 6.**
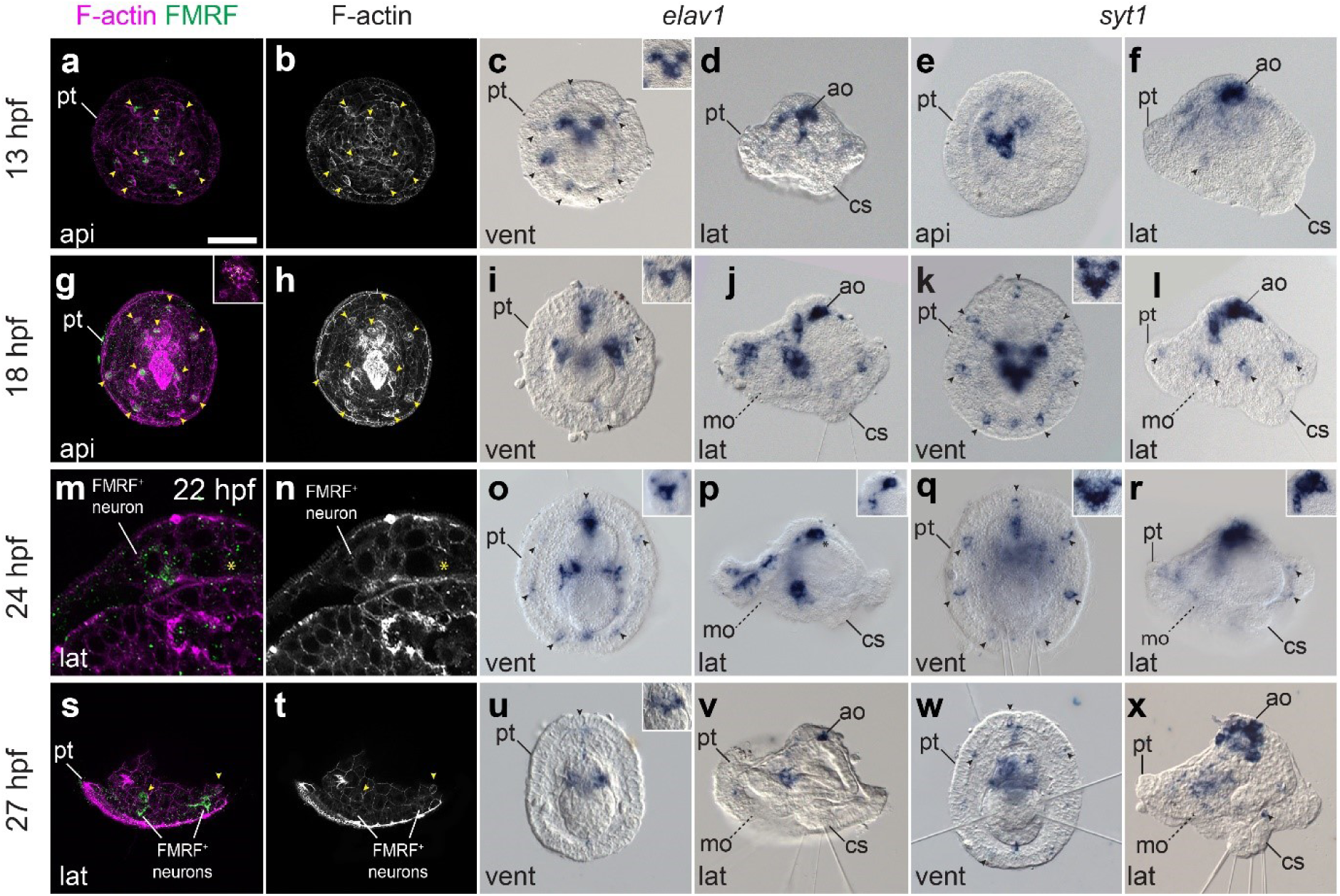
Early neural development during elongation and early mitraria. CLSM images of F-actin and FMRFamide^+^ elements and Differential Interference Contrast (DIC) images showing expression of *elav1* and *syt1* from (**a**) 13 hpf to (**x**) 27 hpf. (**a–b**,**g–h**) There are no FMRFamide^+^ cells except for ten refringent globules (yellow arrowheads) in the apical organ and the prototroch either at 13 or 18 hpf. (**c–d**) A V-shaped apical organ composed of three *elav1^+^* and *syt1*^+^ cells and seven prototrochal cells (black arrowheads) expressing *elav1* are the first neurons to appear. (**i–l**) By 18 hpf *elav1^+^* cells are positioned anterior and lateral to the gut reaching the ventral side of the larva. (**m–n**) The first FMRFamide^+^ cells are present by 22 hpf in the apical organ and by 24 hpf axons extend from the apical organ to an an FMRFamide^+^ prototrochal ring connecting seven FMRFamide^+^ cells in the prototroch (**s–t**) (See Supp Figure 1f–g). (**u–x**) By 27 hpf, more cells express *syt1* in the apical organ, but *elav1* is now mostly restricted to neurons on the ventral side where the juvenile will start forming. an: anus; ao: apical organ; cs: chaetal sac; mo: mouth; mt: metatroch; nt: neurotroch; pt: prototroch. Scale bar is 50 µm.

From 22 to 27 hpf, the embryo matures into a mitraria larva. The circular muscles surrounding the foregut appear at 22 hpf (Fig. 4h–i). The initially sparse ciliated cells in the hyposphere are now more numerous and expand to cover most of this larval region as a secondary ciliary band, except the chaetal sac (Fig. 4h). Following Wilson (Wilson, 1932) and Emlet and Strathmann (Emlet and Strathmann, 1994), we herein refer to this ciliary band as the mitraria metatroch. As the prototroch matures, the mitraria also starts swimming in the water column and can already feed by 22 hpf. After 27 hpf, the mitraria larva is slightly larger, more ciliated (Fig. 4j–k) and have longer and more robust chaetae (Fig. 6).

### Nervous system development in the early mitraria larva

Previous neuroanatomical studies revealed a nervous system in the early mitraria larva consisting of one or two FMRFamide immunoreactive cells in the apical organ and possibly a lateral FMRFamide^+^ axon (Helm et al., 2016; Martín-Durán et al., 2018). However, most other annelid trochophore-like larvae display more complex nervous systems at the moment of hatching (Marlow et al., 2014; Verasztó et al., 2020; Vergara et al., 2017), thus suggesting that the early mitraria larva emerges with a rudimentary nervous system that matures as the larva grows. To test this scenario, we identified and analysed the embryonic expression of the pan-neural marker gene *elav1* (Carrillo-Baltodano et al., 2019; Denes et al., 2007; Koushika et al., 1996; Kumar et al., 2020; Meyer and Seaver, 2009) and the mature neuron marker *synaptotagmin1* (*syt1*) (Carrillo-Baltodano et al., 2019; Kerbl et al., 2016a; Kumar et al., 2020; Meyer et al., 2015; Rizo and Rosenmun, 2008; Santagata et al., 2012; Simionato et al., 2008) (Fig. 6). The earliest stage showing expression of *elav1* and *syt1* is 13 hpf, when *elav1^+^* and *syt1^+^* cells, which we consider neurons, locate around the forming apical tuft, conforming a V-shaped apical organ composed of one large cell in the centre and two smaller lateral neurons (Fig. 7c–f, and insets within). As part of this incipient apical organ, there are two acetylated tubulin^+^ flask-shaped monociliated cells (Fig. 4c,e, Supp. Fig. 2a). In addition, there are three anterior and four posterior neurons by the prototroch expressing *elav1* and *syt1* weakly (Fig. 6c–f). These seven neurons are apparently associated with the seven refringent globules. At this stage, none of the *elav1^+^* and *syt1^+^* cells are immunoreactive against FMRFamide (Fig. 7a) and serotonin (Supp. Fig. 2h). Instead, the refringent globules are positive against FMRFamide and serotonin (yellow arrowheads, Fig. 7a, Supp. Fig. 2f–h), but considering they do not express *elav1* or *syt1*, it is uncertain whether these cells are of neural or neurosecretory nature. By 18 hpf, more *elav1*^+^ and *syt1*^+^ cells appear in the circumesophageal connectives, the ventral tissue where the juvenile rudiment will form, and in a domain anterior to the foregut (Fig. 6i–l). At least two tubulin^+^ cells make the loop of cilia that extrudes out of the larva in the apical tuft (Fig. 4g, Supp. Fig. 2b,e). These cells are in close vicinity to the retractor muscles and the two lateral axons that run from the apical organ towards the ventral side. This general pattern of the nervous system observed at 18 hpf remains in the early mitraria larva, where neurons in the apical organ and in the seven neuronal cells of the prototroch become immunoreactive against FMRFamide, forming a prototrochal ring (Fig. 6m–r, Supp. Fig. 2f–g). However, the expression of *elav1* decays in the 27 hpf mitraria, only remaining in a few cells of the apical organ and in the area where the juvenile rudiment will form (Fig. 6u–x). Altogether, our data demonstrates that neurogenesis starts earlier than previously recognized in *O. fusiformis*, initiating on the apical side and extending posteriorly in conjunction with other morphogenetic events.

**Figure 7.**
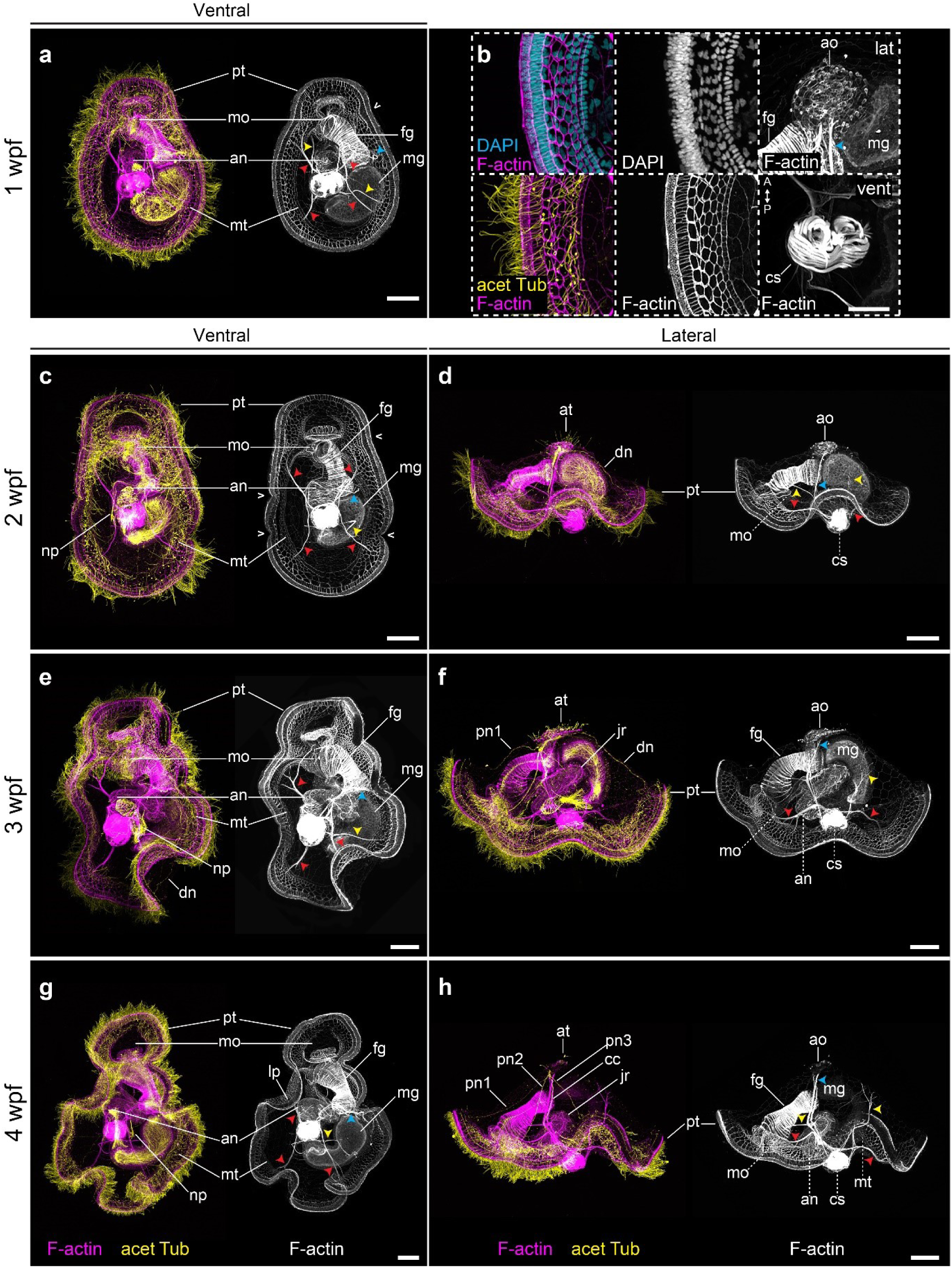
Late mitraria larvae and juvenile rudiment development. CLSM images of late larval stages from (**a**) 1wpf to (**h**) 4wpf. (**a**), (**c**), (**e**) and (**g**) are ventral views with anterior facing up, while (**d**), (**f**) and (**h**) are lateral views with anterior facing left. (**b**) shows close ups of specimens at 1wpf. (**a**) The mature larva continues to grow and develop more musculature to connect the chaetal sac to different parts of the episphere and the hyposphere. The prototroch starts making bends (open arrowheads), and the metatroch is cleared from the chaetal sac and the ventral area where the juvenile rudiment is developing. (**b**) Both the hyposphere and episphere epithelial cells enlarge, the apical organ becomes more prominent and is full of microvilli, and the chaetal sac muscles become more robust. (**c–h**) The apical organ now is connected to the prototrochal ring via peripheral and dorsal nerves. Cyan arrowheads point to the retractor muscles. Yellow arrowheads point to esophageal and dorsal levator muscles. Red arrowheads point to branching muscle connecting the peripheral regions of the ventrolateral hyposhere and the dorsolateral hyposphere muscles. an: anus; ao: apical organ; at: apical tuft; cc: circumesophageal connectives; cs: chaetal sac; dl: dorsal levator; dn: dorsal nerve; fg: foregut; jr: juvenile rudiment; lp: lappet; mg: midgut; mo: mouth; mt: metatroch; np: nephridia; nt: neurotroch; pn1**–**pn3: peripheral nerves 1 to 3; pt: prototroch. Scale bar is 50 µm. In (**b**) the scale bar is 25 µm.

### Growth and competence of the mitraria larva

The 27 hpf mature mitraria is an active planktotrophic stage that acquires competence and undergoes a dramatic metamorphosis after about three weeks at 15 °C. One week post fertilization (wpf), the mitraria grows considerably in size, partly through cell division, but likely also through cell growth and cell shape changes (Fig. 7a–b). The epithelial cells of both the episphere and hyposphere become large and very thin (Helm et al., 2016; Wilson, 1932), often displaying a variety of nuclear shapes and sizes. Related to this growth, the prototroch bends (Fig. 7a, 8a,e) and the metatroch gets spatially restricted and cleared from most of the hyposphere. In both the prototroch and the metatroch of the mitraria, we found evidence of cell proliferation (Fig. 8a,e). Two clearly distinct ciliary bands thus form, leaving a food groove in between (Fig. 7a–b, 8a) (Emlet and Strathmann, 1994; Smart and Von Dassow, 2009; Wilson, 1932). Internally, the apical organ becomes more elaborated and new pairs of muscles appear. In addition to the esophageal, the dorsal levator and retractor muscles (Fig. 7a–b), the chaetal sac connects to the hyposphere through the ventrolateral hyposhere muscles and the dorsolateral hyposphere muscles (Fig. 7a–b). The muscles surrounding the chaetal sac also acquire maturity (Fig. 7b). Already at this stage the juvenile rudiment starts to form, initially as an ectodermal invagination between the mouth and the anus, with the latter eventually becoming incorporated into the newly forming trunk (Fig. 7a, 8a,e).

**Figure 8.**
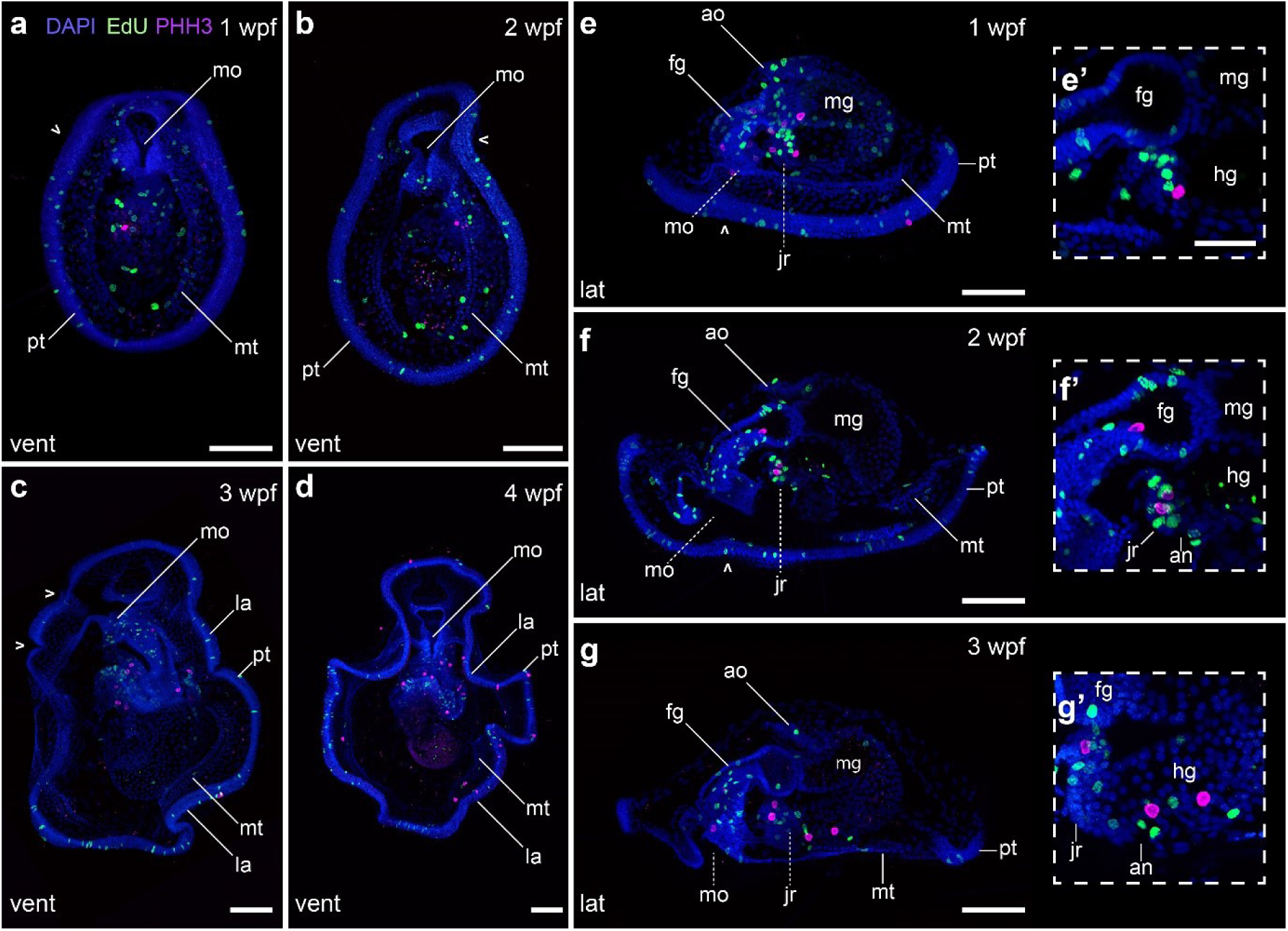
Growth of the late mitraria larvae. CLSM images of EdU and phosphoHistone 3 (PHH3) labeling on mitraria larvae from (**a**), (**e**) 1 wpf to (**d**) 4wpf. Images in (**a–d**) are ventral views with anterior facing up, while those in (**e–g**) are lateral views with anterior facing left. (**e’–g’**) are close up sections of (**e–g**). (**a–g**) Proliferation in the late mitraria continues mostly in the ciliary bands, the gut and the developing juvenile rudiment. Open arrowheads point to the flexions of the ciliary band. (**e’–g’**) The juvenile rudiment has proliferative cells and a terminal mitotic cell, in what it could represent a posterior growth zone. an: anus; ao: apical organ; bp: blastopore; cs: chaetal sac; fg: foregut; hg: hindgut; jr: juvenile rudiment; la: lappet; mg: midgut; mo: mouth; mt: metatroch; nt: neurotroch; pt: prototroch. Scale bar is 50 µm. In (**e’–g’**) the scale bar is 25 µm.

During the following two weeks, the bends of the prototroch exaggerate, forming anterior and posterior lappets (Fig. 7c–h, 8b–d,f–g), a pair of eyes appear near the apical tuft (Helm et al., 2016; Wilson, 1932) and the nephridia enlarge on both left and right sides of the hyposphere (Fig. 7c,e,g) (Gąsiorowski et al., 2020; Smith et al., 1987; Wilson, 1932). As previously described (Helm et al., 2016; Helm et al., 2017), the nervous system also becomes more complex, with new nerves (peripheral nerves 1–3 and dorsal nerve) connecting the apical organ to various regions of the prototroch, and the circumesophageal connectives to the juvenile nerve cord (Fig. 7c–h). At these late mitraria stages, expression of *elav1* is only found in the developing nerve cord of the juvenile rudiment (Fig. 9d–f), suggesting that this tissue is the active site of neurogenesis. Mature FMRFamide^+^ neurons also expressing *syt1* compose the rest of the larval nervous system including the apical organ, which will be the brain of the juvenile (Helm et al., 2016; Wilson, 1932), and the juvenile rudiment (Fig. 9a–c,g–i). An FMRFamide^+^ nerve innervates the chaetal sac (Fig. 9c), which suggests this neuropeptide might control chaetal movements, as in a brachiopod larva (Thiel et al., 2017).

**Figure 9.**
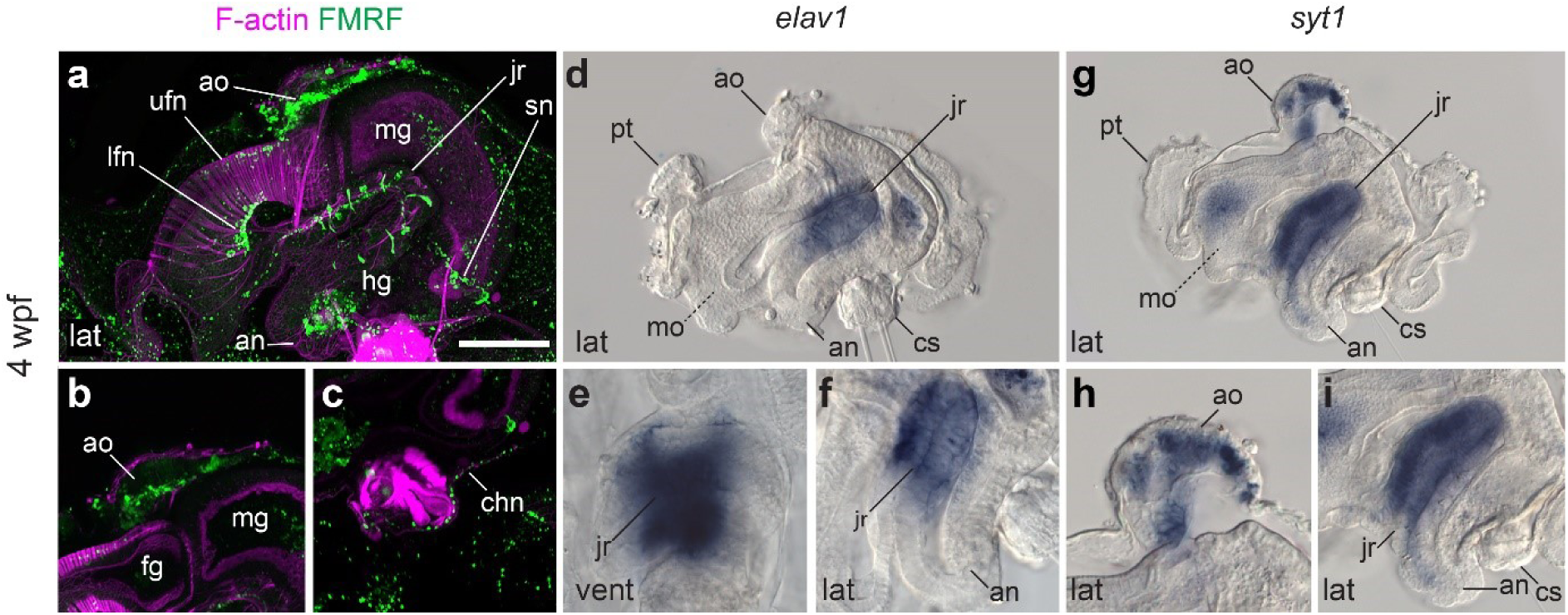
Neural development in the late mitraria larvae. CLSM images of F-actin and FMRFamide^+^ elements and Differential Interference Contrast (DIC) images showing expression of *elav1* and *syt1* at 4wpf. Images are in lateral view, except for (**e**) which is in ventral view. Bottom row are close-up sections of the animals from the top row. (**a–c**) More FMRFamide^+^ neurons and neurites continue to form as the mitraria matures, including those innervating the foregut and the peripheral nerves of the juvenile rudiment (**a**), and the chaetal sac (**c**). See Helm et al. (Helm et al., 2016) for an extended description of the development of the FMRFamidergic and serotonergic nervous system of the late mtiraria. *elav1* expression has declined from the apical organ and is now restricted to the juvenile rudiment (**a–c**), instead, *syt1* is expressed in the mature neurons of the apical organ and the juvenile rudiment. an: anus; ao: apical organ; chn: chaetal sac nerve; cs: chaetal sac; fg: foregut; hg: hindgut; jr: juvenile rudiment; la: lappet; lfn: lower foregut nerve; mg: midgut; mo: mouth; mt: metatroch; nt: neurotroch; pt: prototroch; sn: sphincter nerve; ufn: upper foregut nerve. Scale bar is 50 µm.

Larval competence is acquired at about 3 wpf at 15 °C, when the juvenile rudiment is well-developed and protrudes out of the hyposphere, right anterior to the chaetal sac. During maturation, the juvenile rudiment grows in a posterior to anterior direction (Smart and Von Dassow, 2009; Wilson, 1932) with the trunk cells extending dorsally from the ventral side to wrap the gut and incorporate the larval digestive system into the developing juvenile trunk. Interestingly, the EdU and phospho-Histone H3 signal suggests there is a presumptive proliferation zone at the most posterior part of the trunk (magenta cells in Fig. 8e’—g’), which could represent a posterior growth zone, like in many other annelids (De Rosa et al., 2005; Giani et al., 2011; Rebscher et al., 2007; Seaver et al., 2005). Metamorphosis is often triggered by adding a pinch of mud to the bowl with competent mitraria larvae. During this drastic event, the apical organ fuses with the developing ventral nerve cord and becomes the juvenile brain (Helm et al., 2016; Wilson, 1932), the larval gut is incorporated into the juvenile, which consumes the rest of the larval tissue while shedding the chaetal sac (Smart and Von Dassow, 2009; Wilson, 1932). Soon after metamorphosis, the juvenile will start forming the sand tube where the adult dwells (Fig. 1b).

## Discussion

Our data provide a detailed analysis of the early development and organogenesis of *O. fusiformis*, defining a staging system and a set of morphological landmarks (Fig. 10) that will serve as reference for future developmental and evolutionary studies in this species, and annelids and spiralians generally.

**Figure 10.**
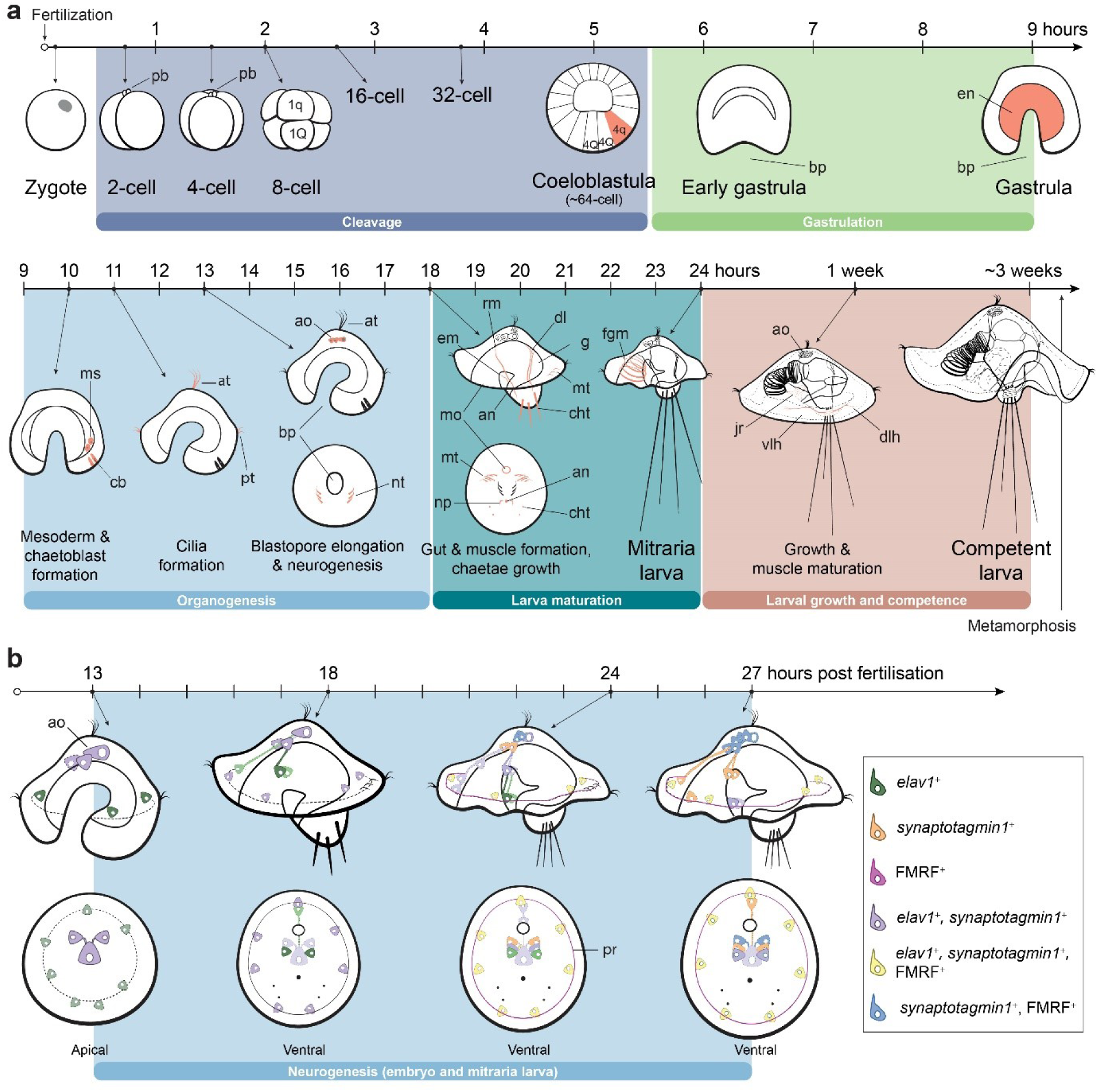
Summary diagram and stage chart of *O. fusiformis* development. (**a**) Time course of the developmental stages from oocyte to late mitraria larvae and (**b**) neurogenesis in the mitraria larvae. Red in (**a**) marks the first appearance of the most relevant cell or tissue at that stage. (**a**) *Top row*: *Owenia fusiformis* undergoes a stereotypical equalspiral cleavage from 0.5 hpf to 5 hpf resulting in the formation of a coeloblastula. Between 5.5 and 6 hpf, the vegetal gastral plate starts gastrulating via invagination, and gastrulation is finished by 9 hpf with the closure of the vegetal blastopore. *Bottom row*: Organogenesis starts at 10 hpf with the formation of the mesodermal precursor cells and chaetoblasts, followed by an apical tuft and an equatorial prototroch at 11 hpf, which divides the embryo in the episphere and the hyposphere. By 13 hpf the blastopore elongates and a neurotroch forms posterior to the blastopore. Myogenesis starts sometime before 18 hpf, when different larval muscles extend from the chaetal sac to the apical organ and the now fully formed gut. Chaetae project out of the chaetal sac and the larvae start swimming. As the larva continues to grow until competence, several other muscles become part of a musculature that connects the chaetal sac with the episphere and hyposphere. (**b**) At 13 hpf, neurogenesis starts from anterior to posterior, with *elav1*^+^ and *syt1*^+^ cells at the apical organ and as the embryo develops, the nervous system differentiate in in a ventral/posterior progression where the nerve cord of the juvenile rudiment will form. FMRFamide^+^ appear by 22 to 24 hpf in the apical organ in the prototroch. an an: anus; ao: apical organ; at: apical tuft; bl: blastocoel; bp: blastopore; cb: chaetoblasts; cht: chaetae; dl: dorsal levator muscles; dlh: dorsolateral hyposphere muscles; em: esophageal muscle; en: endoderm; fgm: foregut muscle; jr: juvenile rudiment; mo: mouth; ms: mesentoblasts; mt: metatroch; np: nephridia; nt: neurotroch; pb: polar bodies; pr: prototrochal ring; rm: retractor muscles; vlh: ventrolateral hyposphere muscles.

### The early spiral cleavage and gastrulation of *Owenia*

Previous descriptions of the early embryonic stages in Oweniids were only available for *O. collaris* (Smart and Von Dassow, 2009). Our study demonstrates that spiral cleavage stages are practically identical between *O. fusiformis* and *O. collaris*, aside from timing differences that are likely due to different culture temperatures (12 °C in *O. collaris* versus 19 °C in *O. fusiformis*). Both *Owenia* species undergo dextral spiral cleavage, and this chirality is already observed as early as in the second cleavage in *O. fusiformis*. Remarkably, the spiral deformation of the actin cytoskeleton in the 2-cell stage embryo has also been described in some mollusk species (Kuroda and Abe, 2020; Shibazaki et al., 2004), and might thus indicate the presence of common early cellular mechanisms underlying embryonic chirality in spiralians. Later on, *Owenia* embryos follow what appears to be a general, probably ancestral pattern for equal spiral cleaving annelids, which includes a symmetric pattern of cell divisions between embryonic quadrants, the formation of a coeloblastula, and gastrulation via invagination (Arenas-Mena, 2007; Costello and Henley, 1971; Freeman and Lundelius, 1992; Gonzales et al., 2007; Mead, 1897; Newby, 1940; Pernet et al., 2015). In some serpulids (Arenas-Mena, 2007; Lambert and Nagy, 2003; Treadwell, 1901), one hesionid (Treadwell, 1901) and one echiuran (Newby, 1940), which are equal cleavers, the asymmetrical division of 2d^2^ into a larger 2d^21^ and smaller 2d^22^ is a landmark of the D-quadrant. However, we did not observe this change in the pattern of cell divisions breaking the embryonic symmetry. Albeit one of the 3q micromeres divides first, we cannot pinpoint which of the four 3q this might be. Therefore, elucidating the quadrant identity of each embryonic region in *O. fusiformis* would require combining cell positional information with potential molecular asymmetries observed in some equal cleaving annelids, such as the activation of the MAPK pathway in the presumptive 4d cell (Lambert and Nagy, 2003).

A remarkable trait of *Owenia* early embryogenesis is the larger size of the 1q micromeres compared to the 1Q macromeres, and subsequently of the 1q^1^ quartet versus their 1q^2^ sisters (Fig. 10a). In the majority of annelids and spiralians, the 1Q macromeres acquire a larger size and a larger allocation of cytoplasm and yolk, because the entire trunk ectoderm, mesoderm and endoderm will form from these lineages (Henry, 2014). However, in annelids with small eggs like *Owenia*, egg size has been correlated with the allocation of cytoplasm to the micromeres and with larval feeding mode (Jones et al., 2016). This reverse ratio between micromere and macromeres is present in another group of annelids, the sipunculans (Boyle and Rice, 2014; Gerould, 1906), where the larger micromere size is thought to contribute to the enlarged amount of apical/anterior ectoderm in the trochophore-like larva of this group, which has big prototrochal cells (Boyle and Rice, 2014). Outside Annelida, enlarged 1q micromeres are common in nemertine embryos, where these first micromere quartet forms the predominant anterior/apical region of their swimming larvae (Henry and Martindale, 1998; Maslakova et al., 2004). Therefore, the presence of relatively larger size of 1q micromeres in *O. fusiformis* could explain the development of a larger episphere in the mitraria larva, as well as the expanded apical/anterior ectoderm in other spiralian larvae.

### Morphogenesis and larval development

Our detailed microscopical study of the mitraria development and growth expands on the descriptions done by Wilson (Wilson, 1932) on specimens collected on the British side of the English Channel (Plymouth, UK) (Fig. 1d). We define the onset of key morphogenetic events, such as chaetogenesis (10 hpf), ciliogenesis (11 hpf), myogenesis and gut formation (in both cases between 13 hpf and 18 hpf) (Fig. 10a). Certain trends become evident when comparing annelids which have a feeding larva like *Owenia* to those which have a non-feeding lecithotrophic larva (i.e., supplied with nutritional yolk by the mother). Feeding larvae tend to form a functional gut earlier (Anderson, 1966; Boyle and Rice, 2018; McDougall et al., 2006; Pernet and McHugh, 2010; Seaver et al., 2005), as well as chaetae and muscle associated with chaetae for defense against predators (McDougall et al., 2006; Pernet and McHugh, 2010; Seaver et al., 2005). On the other hand, non-feeding larvae tend to accelerate the development of juvenile structures (e.g. a definitive brain and muscles) that will be carry over after metamorphosis (Anderson, 1966; Boyle and Rice, 2018; Meyer et al., 2015; Page, 2009; Pernet and McHugh, 2010; Seaver et al., 2005). However, interspecific comparisons of these events are often challenging, because of technical variability in rearing methods and due to species-specific differences in the timing of morphogenesis.

Annelid trochophore larvae have multiple and distinct ciliary structures, including an apical tuft, a prototroch, a neurotroch, a metatroch and/or a telotroch (Nielsen, 2004; Rouse, 1999). Our study demonstrates that the first cilia to appear in *O. fusiformis* are those in the apical tuft and the prototroch, followed by the neurotroch, and lastly the metatroch (Fig 10a). As observed in other annelid studies with enough temporal resolution, the prototroch and apical tuft develop quicker, presumably providing the early larvae with a way to move and sense the environment (Fischer et al., 2010; Kumar et al., 2020; McDougall et al., 2006), while the metatroch differentiates soon after, in preparation for feeding. Unlike the definitive prototrochal cells of other annelid and spiralian larvae (Bird, 2012; Emlet and Strathmann, 1994; Özpolat et al., 2017), ciliated cells in Owenids are monociliated (Gardiner, 1978; Smart and Von Dassow, 2009), and thus can continue dividing once differentiated and integrated in the ciliated band, as previously observed in *O. collaris* (Bird, 2012; Smart and Von Dassow, 2009) and our study shows for *O. fusiformis* (Fig. 5, 8). Indeed, the prototroch putatively arises from a row of equatorial cells that actively divide in the early post-gastrula (Fig. 3h), and continuous proliferation of cells in both the prototroch and the metatroch contribute to expand these bands and elaborate the feeding apparatus of the larva (Fig. 5, 8). This continuous growth and bending of the ciliary bands is similar to what is observed in other larvae with monociliated cells, such as those of deuterostome invertebrates (Bird, 2012), and help the mitraria larvae to be more efficient at food collection and swimming (Emlet and Strathmann, 1994).

Our study demonstrates a previously unrecognized complexity in the nervous system of the early mitraria larva (Fig. 6 and 10b). The expression of neural genes revealed the presence of neurons in the apical organ as early as 13 hpf (Fig. 6), soon after the appearance of the apical tuft at 11 hpf (Fig. 4a–c). Likewise, we found neurons positioned on the apical margin of the prototroch, which presumably control the ciliary beating of the prototrochal band and connect it to the apical organ (Helm et al., 2016). These early neurons do not initially display common neurotransmitters (e.g. serotonin and FMRFamide), but FMRFamide^+^ cells appear in the apical organ soon after 22 hpf, significantly earlier than the seven days post fertilization previously reported (Helm et al., 2016). Neurogenesis (i.e. differentiation of the first neurons and neurites) thus progresses from anterior to posterior in *O. fusiformis*. In some members of Sedentaria (e.g. *Spirobranchus lamarcki* (McDougall et al., 2006) and *Malacoceros fuliginosus* (Kumar et al., 2020)) and Errantia (e.g. *Platynereis dumerilii* (Starunov et al., 2017) and *Phyllodoce maculata* (Voronezhskaya et al., 2003)), neurogenesis starts with a dual anterior and posterior differentiation of pioneer neurons, which has been proposed as the ancestral condition to Pleistoannelida or even Annelida as a whole (Kumar et al., 2020). However, our findings in *O. fusiformis* are in agreement with observations in sipunculans (Carrillo-Baltodano et al., 2019; Kristof and Maiorova, 2015; Kristof et al., 2008) and dinophilids (Kerbl et al., 2016a; Kerbl et al., 2016b), thus suggesting that an initial anterior/apical specification of neurons and pioneer neurons is common in groups outside Pleistoannelida. Interestingly, either condition is present haphazardly across members of Spiralia (Supp Table 1), suggesting that either the presence of two neurogenic centers was ancestral and got repeatedly lost in early annelid lineages and other spiralians, or the development of the nervous system from both anterior and posterior pioneer neurons evolved convergently multiple times in Annelida and Spiralia. Finally, the rapid specification of neurons in the apical organ and near the prototroch in *O. fusiformis*, in combination with the early development of chaetoblasts (within 13hpf) is in line with a selective pressure on the pelagic planktorophic larvae, which needs to have a functional nervous system in order to response to multiple environmental cues in the water column (Carrillo-Baltodano and Meyer, 2017; Conzelmann et al., 2011).

## Conclusions

Our study describes early cleavage, gastrulation and larval development in the palaeoannelid species *O. fusiformis*. This species exhibits a canonical equal spiral cleavage programme, which as in other equal cleaving annelids, appears to be associated with the formation of a coeloblastula, gastrulation via invagination, and the development of a feeding larva (Arenas-Mena, 2007; Costello and Henley, 1971; Freeman and Lundelius, 1992; Gonzales et al., 2007; Mead, 1897; Newby, 1940; Pernet et al., 2015). Given the distribution of these characters across annelid and spiralian phylogeny, these could represent ancestral developmental traits for Annelida. Further studies in magelonids, the sister lineage to oweniids, will clarify whether all palaeoannelids exhibit similar developmental strategies. In addition, our study sets a timing for the onset of major morphogenetic and differentiation events during organogenesis, uncovering that neurogenesis starts soon after gastrulation, much earlier than previously recognized. Altogether, our study establishes a reference description and staging of the embryogenesis of *O. fusiformis*, an emergent research annelid species for cell, developmental, and evolutionary biology.

## Methods

### Adult culture, spawning and in vitro fertilization

Adults of *O. fusiformis* Delle Chiaje, 1844 (Fig. 1b–c) and mud were collected and shipped from the subtidal waters near Roscoff, France (Fig. 1d). The worms were acclimated inside aquaria with aerated artificial sea water (ASW) and mud at 15°C for about one week after arrival. On the day of the artificial fertilization, a group of individuals were placed in a glass bowl with clean ASW, and later relaxed with 8% MgCl_2_ for 10-12 minutes (Supp Fig. 2a–b). Juveniles and adults of *O. fusiformis* build tubes out of sand (Fig. 1b). Therefore, after relaxation each tube was opened manually, and then the animals were placed individually in a well of a 6-well plate containing 0.22 µm filtered-ASW (FSW) (Supp Fig. 2c). The sex of each animal was determined by protruding a small opening of the coelomic cavity with fine forceps, allowing for either oocytes or sperm to be observed. There is a 50:50 ratio of males and females (Supp Fig 2e). All males were combined into a single bowl with enough water to keep them wet. On the other hand, groups of 4-5 females were placed into separate bowls (Supp Fig 2d). Females were dissected to release the oocytes inside the coelom. The oocytes were passed through a 150 µm mesh to remove remaining pieces of female tissue and subsequently through a 70 µm mesh to filter small immature oocytes and coelomic content out. During adult dissection, both male and oocyte bowls were kept at 4°C to prevent premature sperm and egg activation, respectively. Oocytes were then incubated at 19°C for ∼45 min to allow germinal vesicle breakdown and thus oocyte activation. Simultaneously, 4-5 males were placed in a glass bowl with a small volume of ASW and dissected to obtain sperm. Sperm activation was monitored by taken one drop of concentrated sperm and observing it under a 40x objective of a compound microscope. The remaining concentrated sperm was diluted into 50 ml of FSW and kept at 19°C. Once the eggs were activated, 125 µl of diluted sperm was pipetted into the glass bowls containing the oocytes. After 30 min, the sperm was washed out with three rinses in FSW, and the embryos were cultured at 19°C until they reach the desired developmental stage. Swimming mitraria emerged at approximately 24 hours post fertilization, when they were transferred to plastic beakers containing 600 ml of ASW and placed on a rocker at 30 rpm in a controlled temperature room at 15°C (Supp Fig. 2f). Three times a week, we exchanged the ASW of the beakers and fed the larvae with 100 µl of Reef Juice – Live Phytoplankton Blend (Reefphyto).

### Proliferation assays

Embryos and larvae from 9 hpf to 4 wpf were incubated in 3 µM 5-Ethynyl-2′-deoxyuridine (EdU; ThermoFisher Scientific, cat#: C10639) in FSW for 15-30 min. Specimens were then relaxed in 8% MgCl_2_ for 10 min and fixed with 4% paraformaldehyde for 1h at RT with constant rocking. EdU incorporation was visualized with a Click-iT EdU Alexa Fluor 594 (ThermoFisher Scientific, cat#: C10639) following the manufacturer’s instructions. Immunostaining (below) was performed after the Click-iT reaction.

### Immunohistochemistry

Antibody staining was carried out as previously described (Helm et al., 2016; Martín-Durán et al., 2016a). Briefly, fixed samples were washed quickly with PBS, permeabilized with washes of PBS+0.1% Triton X-100 + 1% bovine serum albumin (PTx+BSA) over 1h, and blocked in PTx + 0.5% normal goat serum (NGS) for 1h. Samples were incubated in primary antibodies diluted in NGS overnight at 4°C while rocking, and subsequently washed four times in 30 min intervals with PTx+BSA while rocking. Secondary antibodies were incubated overnight diluted in NGS and washed four times in 30 min intervals with PTx+BSA. Before imaging, samples were washed with PBS and cleared with 70% glycerol in PBS. Primary antibodies are as follow: 1:1000 rabbit anti-phospho histone H3 (Cell Signaling Technology, cat#: 9701S), 1:800 mouse anti-acetylated tubulin (clone 6-11B-1, Millipore-Sigma, cat#: MABT868), 1:600 rabbit anti-FMRFamide (Immunostar, cat#: 20091). Secondary antibodies are as follows: 1:800 goat anti-mouse AlexaFluor 488 (ThermoFisher Scientific, cat#: A32731), 1:800 goat anti-mouse AlexaFluor 647 (ThermoFisher Scientific, cat#: A-21235) and 1:800 anti-rabbit AlexaFluor 555 (ThermoFisher Scientific, cat#: A-21428). To visualize nuclear and F-actin staining, animals were incubated in 5 µg/ml DAPI (ThermoFisher Scientific, cat#: D3571) and 1:100 AlexaFluor 488 Phalloidin (ThermoFisher Scientific, cat#: A12379) overnight at 4°C in combination with the secondary antibodies.

### Gene isolation, riboprobe synthesis and in situ hybridization

*Owenia fusiformis* orthologs of *elav1* and *syt1* were mined from a published transcriptome (SRA# SRX512807). Fragments of the coding sequences of these genes were amplified to use as templates for RNA probes, using a nested PCR with a T7 universal primer. Gene-specific primers with a T7 adapter on the 5’ end of the reverse primer (underlined) were used as follows: *elav1* 587 bp, forward primer 5’-CCAACAACAGGGCTATCTAAAGG and reverse primer 5’-<u>GCCCCGGC</u>GGAGACTTGCGATTACTGGT; *syt1* 1002 bp, forward primer 5’-AGGGATAGTGGCCGTTCTAC and reverse primer 5’-<u>CCCCGGC</u>CGTAACTCTGTACCCGATGC. Animals for *in situ* hybridization (ISH) were washed from the fixative with PBS + 0.1 Tween-20 (PTw), dehydrated in a gradient of methanol and stored at −20°C in 100% methanol. ISH protocol was carried out as described elsewhere (Martín-Durán et al., 2016a), except that animals were permeabilized with 5 µg/ml Proteinase-K for 90 seconds (ThermoFisher Scientific, cat# AM2548), and hybridized with 1ng/µl of RNA probe for a minimum of 48 hours at 62°C.

### Orthology assignment

Multiple protein alignments (MPA) were constructed with MAFFT v.7 (Katoh et al., 2018) using a L-INS-i method. Poorly aligned regions were removed with gBlocks (Talavera and Castresana, 2007). Maximum likelihood trees were constructed with RAxML v.8.2.11 (Stamatakis, 2014) using an LG+G+F model and visualized with FigTree (https://github.com/rambaut/figtree/) (Supp Fig. 3). Accession numbers for the sequences included in the MPAs are listed in Supp Table 2.

### Imaging processing

Differential Interface Contrast (DIC) images were taken with a Leica CTRMIC compound scope couple with a Infinity5 camera (Lumenera). Confocal laser scanning microscopy (CLSM) images were taken with a Leica SP5. CLSM Z-stack projections were built with ImageJ2 (Rueden et al., 2017). To reduce some of the background noise at early stages, the DAPI and phalloidin channels were multiplied and square rooted. This product was subtracted from the original images, as previously described (Smart and Von Dassow, 2009). Autofluorescent dust and precipitates near the animals were digitally removed using Photoshop CC version 14.0 (Adobe Systems, Inc.). DIC images were digitally stacked with Helicon Focus 7 (HeliconSoft). Brightness and contrast were edited with Adobe Photoshop and figures built with Adobe Illustrator CC (Adobe Systems, Inc.).

## Declarations

### Authors’ contributions

AMCB and JMMD conceived the study. AMCB, OS and JMMD collected and fixed the specimens. AMCB and KG performed the immunostaining. AMCB cloned and performed phylogenetic analyses and AMCB and JMMD performed expression analyses. AMCB obtained all images. AMCB and JMMD wrote the manuscript. All authors contributed to the interpretation of the data.

## Acknowledgements

We thank the staff at Station Biologique de Roscoff for their help with collections and animal supplies and all members of the Martín-Durán lab for support and valuable comments on the manuscript.

---

**Supplementary Table 1.** First appearance of neurons along the anterior(apical)-posterior axis in different groups of Spiralia. [csv file]

**Supplementary Table 2.** Accession numbers of protein sequences used for the alignments of Elav and Synaptotagmin proteins. [csv file]

**Supplementary figure 1.**
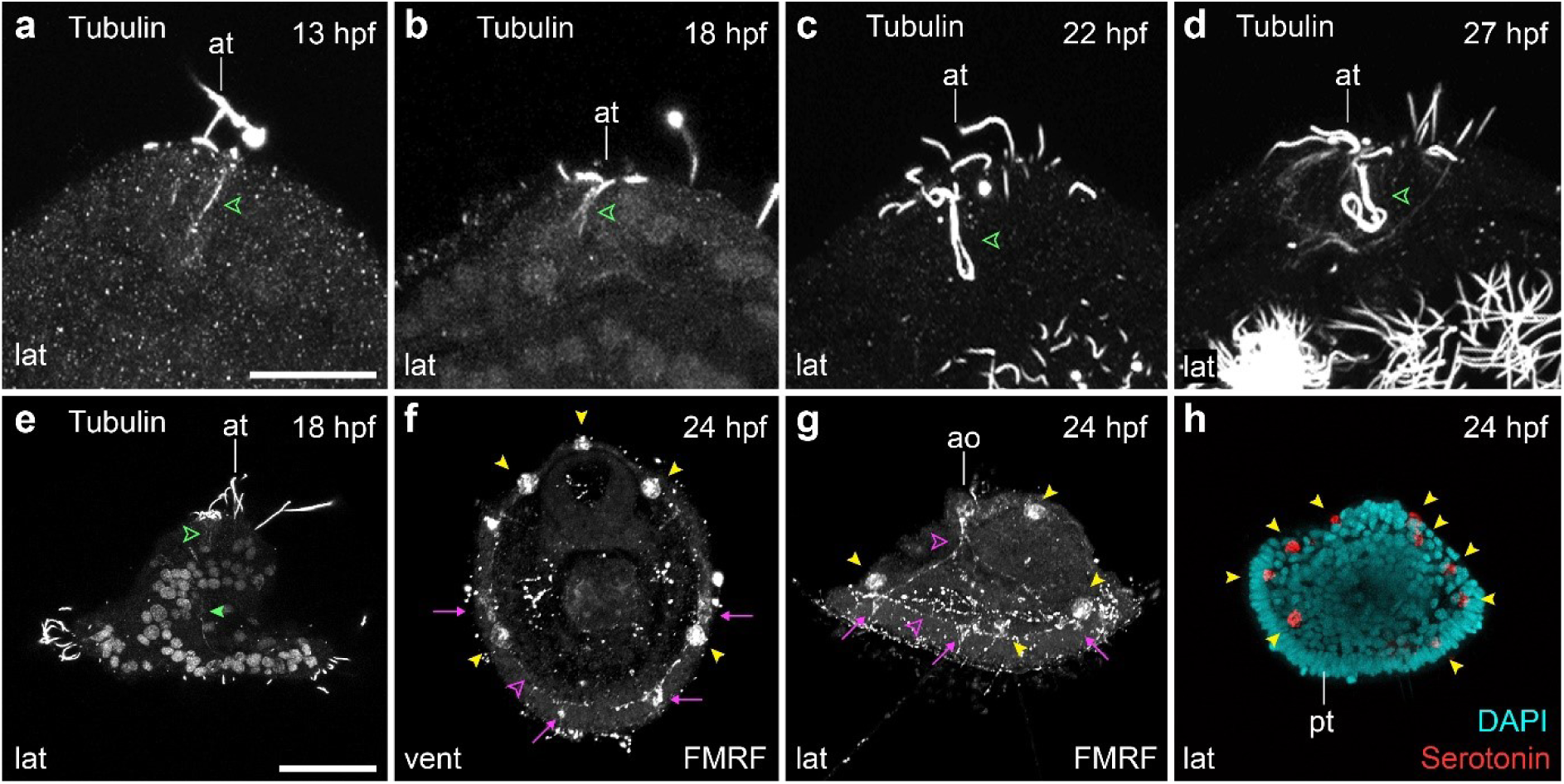
Tubulin^+^ and FMRFamide^+^ elements during neurogenesis. CLSM images early larval stages from (**a**) 13 hpf to (**h**) 24 hpf. (**a–d**) Close ups of the apical tuft innervation into the apical organ (green open arrowhead). (**e**) By 18 hpf an axon connects the apical organ with the prototroch (green closed arrowhead). (**f–g**) The apical organ connects with the neurons along the prototroch (magenta arrows) start to be FMRFamide immunoreactive by 22–24 hpf (see Fig. 6m–n), the latter forming an FMRFamide^+^ prototrochal ring (magenta arrowheads) which connects with the apical organ via an FMRFamide^+^ axon (magenta arrowheads). (**g–h**) The refringent globules that develop from 11 hpf, by 24 hpf are both FMRFamide^+^ and serotonin^+^. ao: apical organ; at: apical tuft; pt: prototroch. Scale is 25 µm in (**a–d**). Scale bar in (**e–h**) is 50 µm.

**Supplementary figure 2.**
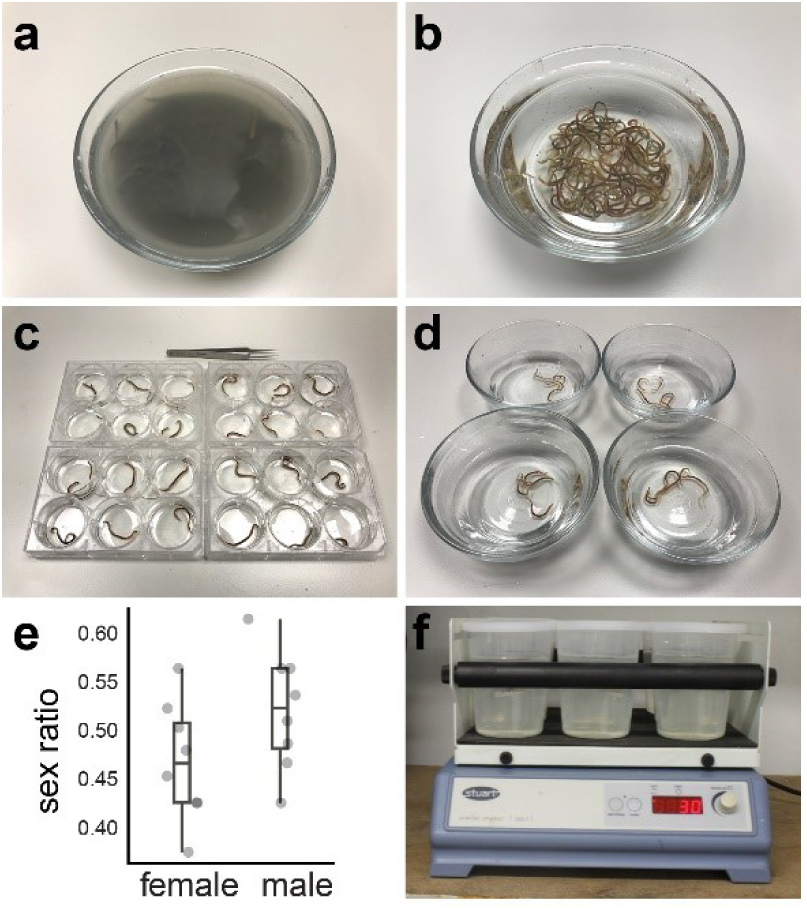
Culture and artificial fertilization. Adults are (**a**) relaxed with MgCl_2_ and (**b**) stripped from their tubes. (**c**) Adults are sorted and and (**d**) separated by sex. (**e**) The sex ratio of adults used in this study were kept very similar. (**f**) After 27 hpf, the larvae were transferred to 600 ml plastic Beakers and grown at 15°C.

**Supplementary figure 3.**
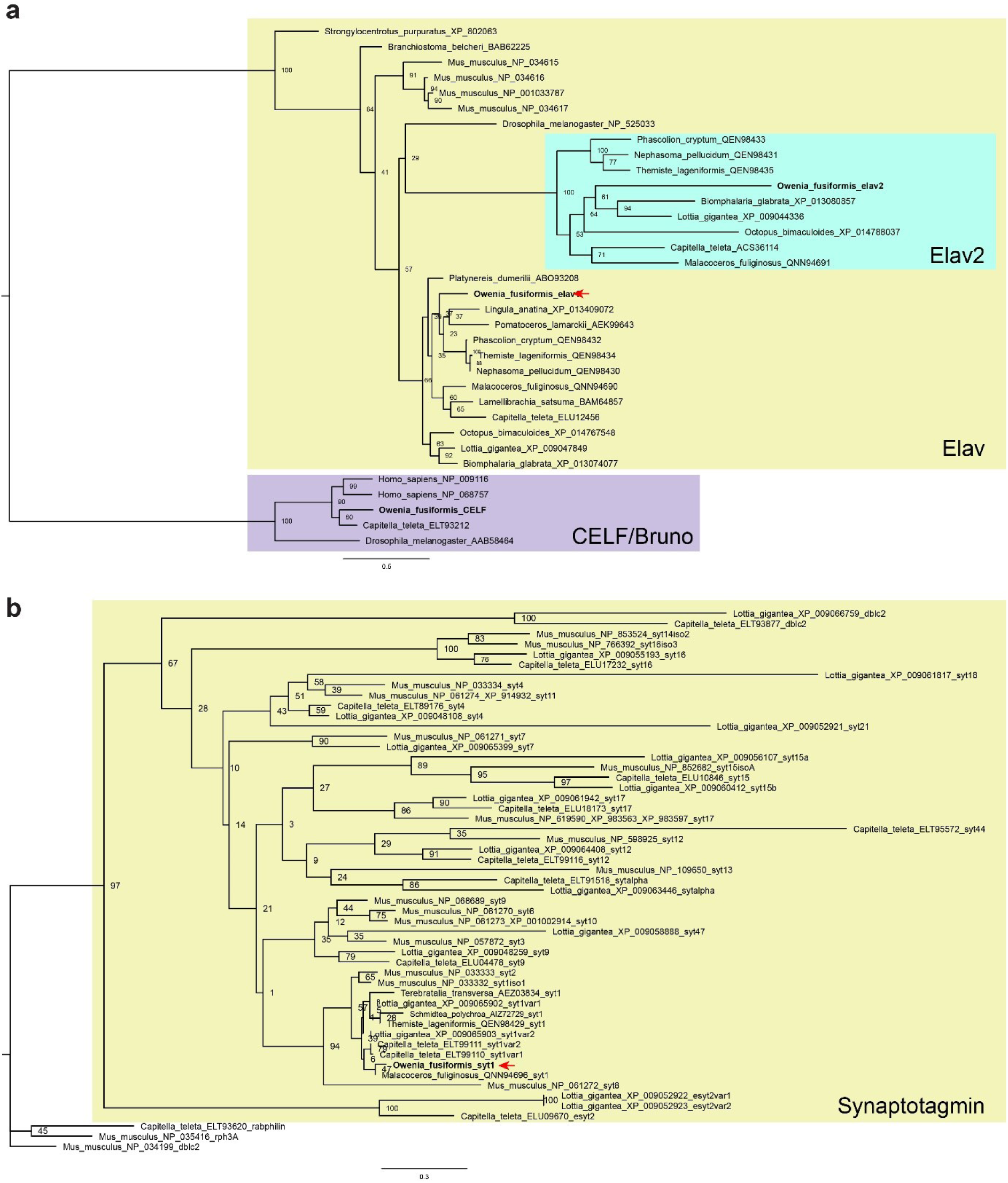
Phylogenetic relationships of *O. fusiformis* Elav1 and Synaptotagmin1 proteins. (**a**) RaxML phylogenetic tree of Elav and (**b**) Synaptotagmin1 of *O. fusiformis*. (**a**) Similar to other spiralians, there are two Elav proteins in *O. fusiformis*. (**b**) *O. fusiformis* Synaptotagmin1 clusters with the Synaptotagmin1 clade from other spiralians. Refer to the methods for the specifics of the phylogenetic analyses and Supp. Table 2 for the accession numbers used.

